# Single-particle cryo-EM analysis of the shell architecture and internal organization of an intact α-carboxysome

**DOI:** 10.1101/2022.02.18.481072

**Authors:** Sasha L. Evans, Monsour M. J. Al-Hazeem, Daniel Mann, Nicolas Smetacek, Andrew J. Beavil, Yaqi Sun, Taiyu Chen, Gregory F. Dykes, Lu-Ning Liu, Julien R. C. Bergeron

## Abstract

Carboxysomes are proteaceous bacterial microcompartments (BMCs) that sequester the key enzymes for carbon fixation in cyanobacteria and some proteobacteria. They consist of a virus-like icosahedral shell, encapsulating carbonic anhydrase and ribulose 1,5-bisphosphate carboxylase/oxygenase (RuBisCO), which catalyses the dehydration of bicarbonate into CO_2_, the first step of the Calvin–Benson–Bassham cycle. Despite their significance in carbon fixation and great bioengineering potentials, the structural characterization of native carboxysomes, including the shell and the internal organization, is currently limited to low-resolution tomography studies. Notably, the degree of heterogeneity of the shell, and the internal arrangement of enzymes, remain poorly understood. Here, we report the structural characterization of a native α-carboxysome from a marine cyanobacterium by single-particle cryo-EM. We determine the structure of RuBisCO enzyme at 2.9 Å resolution. In addition, we obtain low-resolution maps of the icosahedral protein shell and the concentric interior organisation. In combination with artificial intelligence (AI)-driven modelling approaches, we exploited these maps to propose a complete atomic model of an intact carboxysome. This study provides insight into carboxysome structure and protein-protein interactions involved in carboxysome assembly. Advanced knowledge about carboxysome architecture and structural plasticity is critical for not only a better understanding of biological carbon fixation mechanism but also repurposing carboxysomes in synthetic biology for biotechnological applications.

## Introduction

Within cells, proteins tend to self-assemble and interact with other proteins or molecules to form active macromolecular machines, such as metabolic organelles ^1–3^, that play a central role in cellular processes ^4,5^. Understanding the precise structures of protein assemblies is imperative for fundamental investigations of the biosynthesis and function of natural protein assemblies and engineering of artificial assembling nanomaterials for new functions ^6^.

Bacterial microcompartments (BMCs) are large macromolecular assemblies widespread in the bacterial kingdom ^7,8^. Unlike their eukaryotic counterparts, BMCs serve as metabolic organelles that have no lipid bilayer and are composed entirely of proteins ^8^. By segregating metabolic enzymes from the cytosol using virus-like protein shells, BMCs are thought to protect the cell from toxic intermediate metabolites and unwanted side reactions, and play pivotal roles in several enzymatic pathways, including autotrophic CO_2_ fixation, organic compound synthesis, catabolic processes, and iron homeostasis ^9–13^.

Despite their diverse range of functions, all BMCs possess a similar overall organization. They consist of a polyhedral proteinaceous shell, reminiscent of viral capsids. This shell encapsulates the enzymes involved in the corresponding metabolic pathway, and acts as a semi-permeable physical barrier for molecule diffusion ^1,14,15^. Structural studies of multiple BMC shell proteins in isolation have shown that they belong to three distinct families: hexamers and pseudohexameric trimers that tile the majority of the shell facets, and pentamers that cap the vertices of the polyhedral shell ^2,16–18^ Although our knowledge about the entire architectures of BMCs is still primitive, high-resolution cryo-EM structures of synthetic BMC mini-shells have provided insight into the organisation of shell proteins and the dynamic nature of these proteinaceous shells for facilitating metabolite entry and exit ^13,19–21^. Carboxysomes were the first BMCs to be discovered ^22^. They are found in cyanobacteria and some chemoautotrophs, and play a key role in carbon fixation ^23^. Carboxysomes contain the enzymes ribulose-1,5-bisphosphate carboxylase/oxygenase (RuBisCO) and carbonic anhydrase (CA). CA catalyzes the conversion of cytosolic bicarbonate (HCO_3_^-^) into CO_2,_ which is subsequently utilised by RuBisCO and fixed onto the 5-carbon sugar ribulose-1,5-bisphosphate (RuBP) as the first step in the Calvin-Benson-Bessham (CBB) cycle^24^. By generating carboxysomes to sequester these enzymes and allow the accumulation of HCO_3_^-^ /CO_2_, bacterial cells can provide an elevated level of CO_2_ around RuBisCO to enhance carbon fixation and overcome the competitive inhibition of RuBisCO carboxylation by O_2_^23,25^. These intrinsic structural features allow carboxysomes to make a significant contribution to the global carbon fixation ^23^. Notably, repurposing carboxysomes is an emerging discipline with applications in crop engineering, metabolic enhancement, bioenergy production, and therapeutics ^23,26–28^.

Carboxysomes can be classified into two distinct groups: α-carboxysomes, encoded for by the *cso* operon, and β-carboxysomes, encoded for by the *ccm* operon ^29^. These two groups are distinctive by their protein composition and the types of RuBisCO encapsulated, belonging to form 1A and form 1B RuBisCO, respectively. Despite having been suggested to be evolved independently from each other to adapt to different ecological niches, these two forms of RuBisCO demonstrate similar affinities for their substrates ^30^. Unlike RuBisCO and carboxysome shell proteins, the CA enzyme is evolutionarily distinct between α-carboxysomes and β-carboxysomes, with distinct structural folds, although they are essential for function and both carry out similar roles ^31^.

RuBisCO is a hexadecameric complex, comprised of eight large subunits and eight small subunits. The structures of RuBisCO from various cyanobacteria and plant species have been solved ^32–36^. In β-carboxysomes, RuBisCO enzymes appear densely organised and form paracrystalline arrays that are important for β-carboxysome biogenesis ^37–40^. In contrast, RuBisCO enzymes have been postulated to assemble concomitantly with the shell during α-carboxysome biogenesis, a process promoted by the intrinsically disordered protein CsoS2 that promote the association between shell proteins and interiors ^18,41^. The organisation of RuBisCO inside the α-carboxysomes is poorly understood. Previous cryo-electron tomography analysis of α-carboxysomes from the chemoautotrophic bacterium *Halothiobacillus* (*H*.) *neapolitanus* and the cyanobacterial strains *Prochlorococcus marinus* MED4, *Synechococcus* sp. WH8102 and WH8109 showed that the RuBisCO and CA enzymes appear to be packed densely and arranged into concentric layers ^42–45^. However, no model has been proposed for the protein arrangement and interactions within the carboxysome and the architecture of the carboxysome shell. High-throughput imaging using X-ray laser outlined the icosahedral shape of the α-carboxysomes from *H. neapolitanus*, but no high-resolution structures of the intact carboxysome and interior organisation were reported ^46^. These studies, together with those of other BMCs, have highlighted the challenges associated with structural characterisation of these large heterogeneous macromolecular assemblies, specifically with great variations in the stoichiometric composition and interactions of individual building components that are adaptive to environmental changes ^21,47,48^.

Here, we report the first single particle Cryo-EM analysis of an intact α-carboxysome from a marine α-cyanobacterium *Cyanobium* sp. PCC 7001 (hereafter *Cyanobium*). We report the structure of its RuBisCO enzyme to 2.9 Å resolution, with the densities present for both the substrate RuBP and attached ligand. In the α-carboxysome, RuBisCO enzymes form four concentric layers encapsulated by the single-layer protein shell. We also report a low-resolution structure of the icosahedral shell, demonstrating a range of dimensions which precludes high-resolution analysis but nonetheless allows us to propose a hybrid structural model for the α-carboxysome shell architecture. Moreover, 3D reconstruction combined with modelling allows us to propose a model for the arrangement of RuBisCO enzymes within the α-carboxysome. The study provides insight into α-carboxysomes assembly, which will inform rational design and engineering of BMC-based nanostructures for diverse purposes.

## Results

### Purification and Single Particle analysis of α-carboxysomes from *Cyanobium* sp. PCC 7001

The *Cyanobium* α-carboxysome proteins are encoded by a 9-gene operon including 5 genes encoding shell proteins (*csoS1D, csoS1A, csoS4A, csoS4B, and csoS1E*), 3 genes encoding cargo enzymes RuBisCO (*cbbL* and *cbbS*) and CA (*csoSCA*), and one gene encoding the scaffolding protein CsoS2 (*csoS2*) (Figure 1a). CsoS1A and CsoS1E contain one Pfam00936 domain, homologous to the prototypical BMC shell hexamer, that tile the majority of the α-carboxysome shell. CsoS1D, containing two fused Pfam00936 domains, shows similarity to pseudohexamer trimers, presumably responsible for passage of large molecules in and out of the carboxysome. CsoS4A and CsoS4B have one Pfam03319 domain and belong to the family of BMC shell pentamers that cap the vertices of the polyhedral shell (Figure 1b).

**Figure 1:**
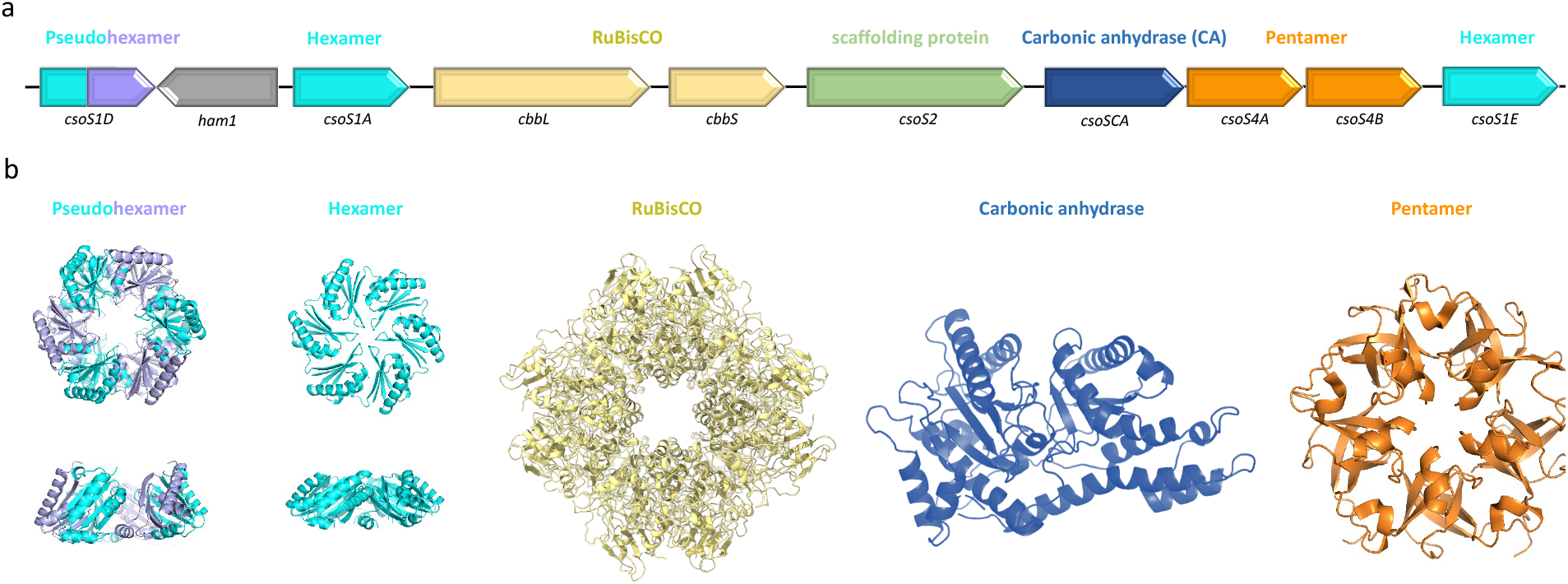
The *Cyanobium* sp. PCC7001 α-carboxysome. **(a)** Gene organisation of the α-carboxysome operon, including genes encoding the shell hexamers (cyan) and pentamers (orange), the scaffolding protein (green), and the cargo enzymes RuBisCO (yellow) and CA (blue). **(b)** Structural model of the corresponding α-carboxysome building proteins, based on the previously-determined structures of homologous proteins.

To isolate functional carboxysomes, we grew *Cyanobium* photosynthetically in BG-11 freshwater medium until the late exponential phase (Figure S1a). Native α-carboxysomes were isolated from *Cyanobium* using sucrose gradient ultracentrifugation and were enriched at the 30-40% sucrose fraction (Figure S1b). SDS-PAGE (Figure 2a) and immunoblot analysis (Figure 2b) of the 40-50% fraction demonstrated the presence of major α-carboxysome components CbbL, CsoS2 and CsoS1A. Mass spectrometry analysis further indicated that the isolated α-carboxysomes comprise all 9 building proteins (Table S1). Among them, RuBisCO subunits (CbbL and CbbS), CsoS2, and CsoS1A are highly abundant proteins, while CsoS4A, CsoS4B, and CsoS1D have low abundance in the α-carboxysome, in good agreement with the mass spectrometry data of α-carboxysomes from *H. neapolitanus* ^55^. Negative-stain EM showed that the isolated α-carboxysomes have a canonical polyhedral intact BMC shape, with an average diameter of ~120 nm (Figure S1c), comparable with previous observations ^30,49^. The ^14^C-based assays of RuBisCO activity confirmed that the isolated α-carboxysomes are catalytically active for carbon fixation (Figure S1d).

**Figure 2:**
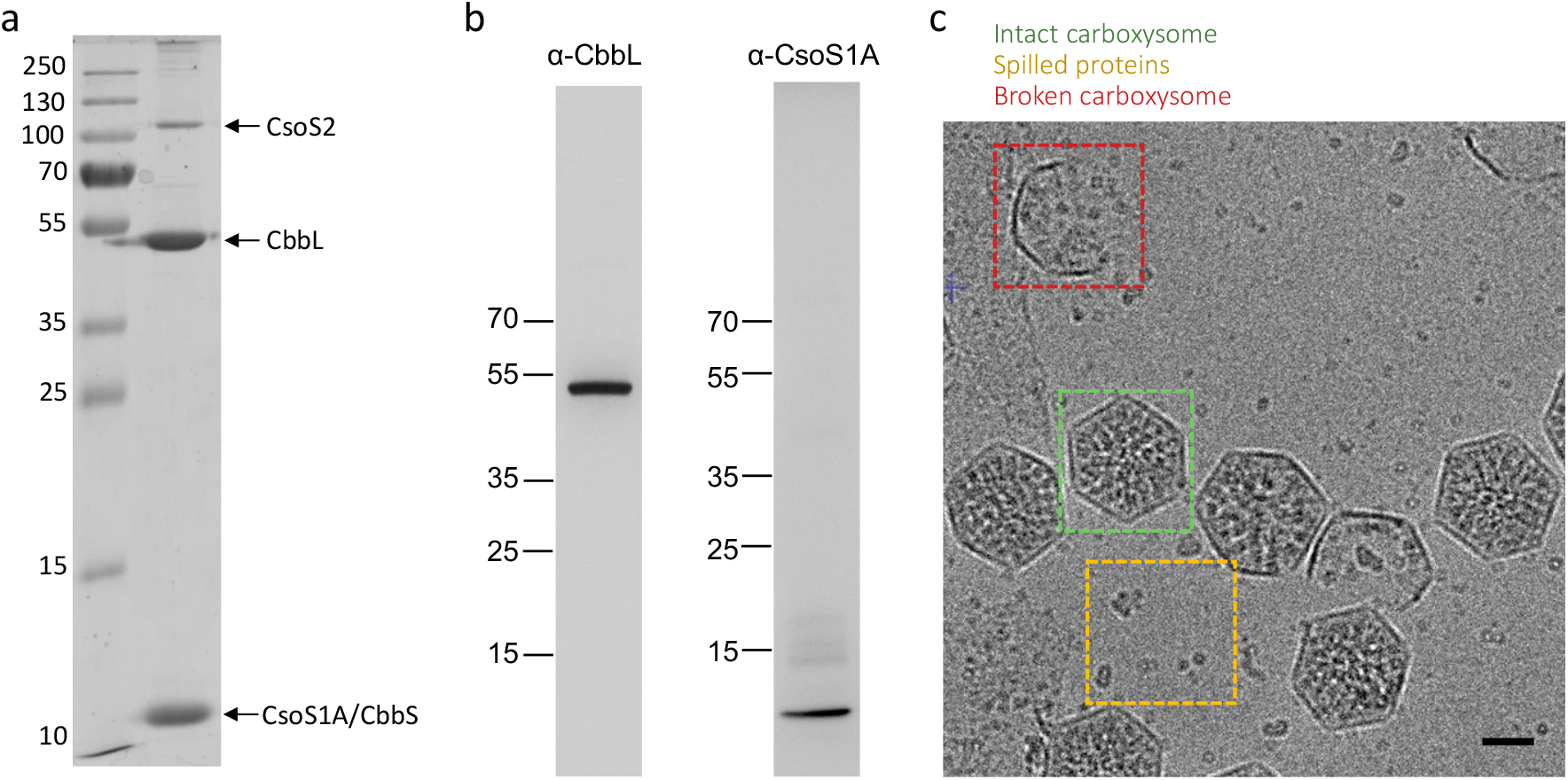
Purification and cryo-EM analysis of the *Cyanobium* α-carboxysome. **(a)** SDS-page of the purified carboxysome. Bands for proteins CsoS2, CbbL, CbbS and CsoS1A could be identified. **(b)** Western blotting of the purified α-carboxysome complex, using antibodies raised against peptides from CbbL and CsoS1, confirming the presence of both proteins. **(c)** Cryo-electron micrograph of frozen-hydrated α-carboxysome samples. Intact BMCs, with incorporated proteins, are visible (green box), along with broken ones (red box). Smaller protein complexes, presumably spilled from these, are also visible (Yellow box). Scale bar: 50 nm.

These intact, functional α-carboxysomes were then subject to single particle cryo-EM to study the three-dimensional architecture of the obtained α-carboxysome assemblies. Initial screening showed a heterogeneous sample containing intact carboxysomes with proteinaceous content, broken carboxysome shell fragments without any cargos inside, and disassembled proteins outside of the carboxysomes (Figure 2c). This deviates from negative-staining EM results (Figure S1c). We postulate that the broken shells largely result from sample handling and/or freezing, and that the disassembled proteins correspond to the proteins that spilled from the broken carboxysomes, including the enzymes RuBisCO and CA, as well as isolated shell components.

### Structure of RuBisCO from native α-carboxysomes

In order to gain structural insights into the α-carboxysomes, we collected a cryo-EM dataset of the sample described above, using a standard, high-magnification (~1 Å/pix^2^) data collection approach. Because of the size of the complex, and its propensity to break (see above), we only obtained few intact carboxysomes fully visible within a micrograph in this dataset. This largely precluded any analysis of the carboxysome complex. However, the spilled particles were readily visible on ice, and we were able to pick these, leading to a set of ~ 3,000,000 particles.

Following initial two-dimensional classifications, clear classes of two distinct molecular species could be identified. Specifically, several classes showed clear 4-fold symmetry, and were visually identified as the RuBisCO (CbbL_8_-CbbS_8_) holoenzyme (Figure S3a). Additional classes were obtained for smaller protein(s), were featureless, and could not be identified based on 2D classes (Figure S3b). We hypothesize that these proteins correspond to a mixture of CA and shell proteins; however, this would require further validation.

We next conducted three-dimensional refinement in the set of particles that could be identified as RuBisCO in the 2D classes. This yielded a 2.9 Å resolution coulomb potential map (Figure 3a, S2c, S2e, Table S2), with eight large subunits (CbbL) and eight small subunits (CbbS) readily identifiable. Using this map, we were able to build an atomic model of the *Cyanobium* RuBisCO enzyme (Figure 3b, Table S2). Notably, density is present in the active site, in a position suitable to be the substrate RuBP (Figure 3c). This observation demonstrated that most RuBisCO enzymes within the carboxysome are active and bound to the substrates, in agreement with the RuBisCO assay results (Figure S1d).

**Figure 3:**
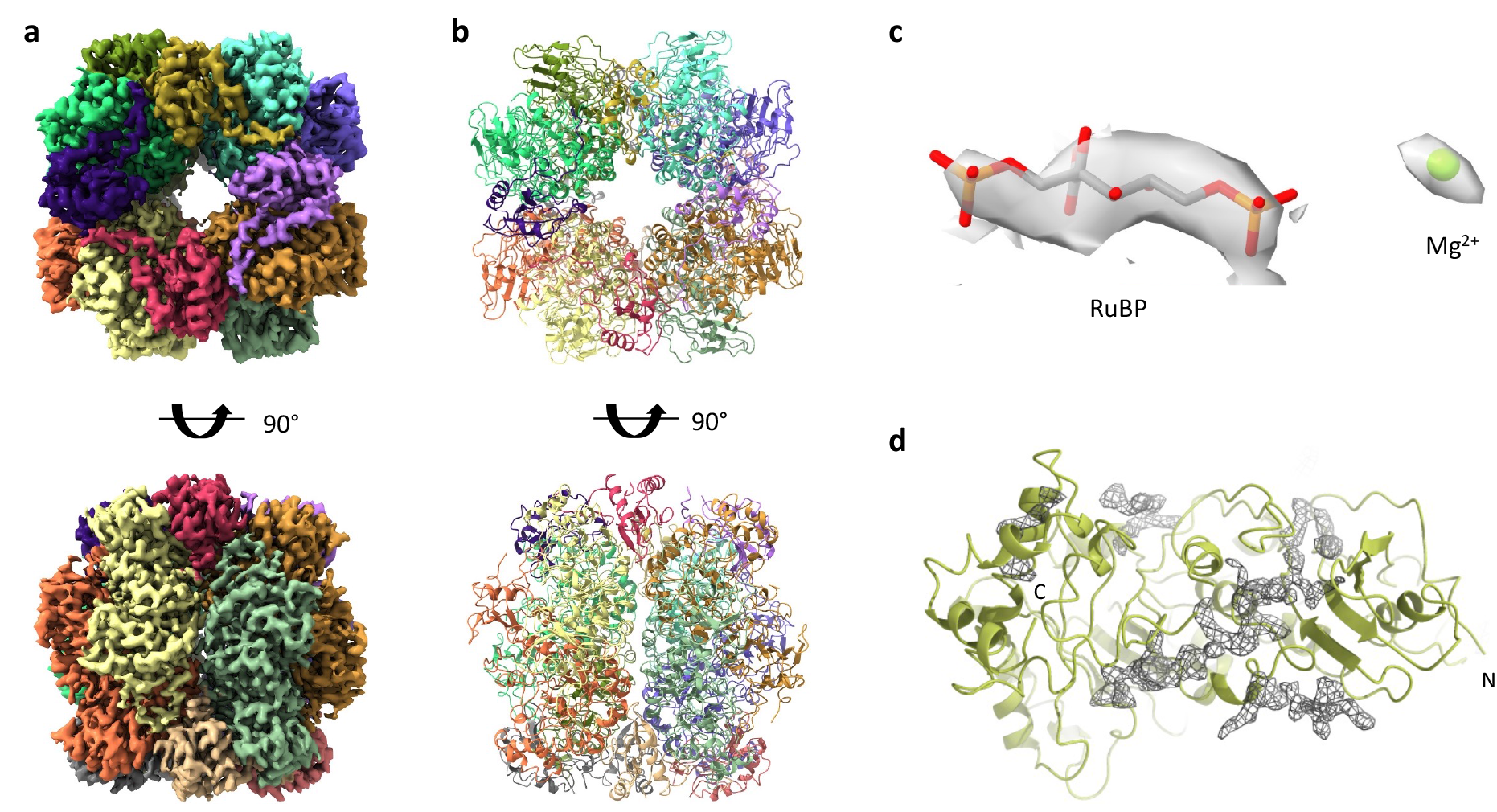
Cryo-EM structure of RuBisCo isolated from 7001 α-carboxysomes. **(a)** Electron potential map of RuBisCO, obtained from the particles of spilled proteins, and segmented by chain. **(b)** Atomic model of the *Cyanobium* RuBisCO enzyme, colored as in (a). **(c)** Close-up view of the active site from one CbbL chain. There is clear density for a ligand, most likely the substrate ribulose-1,5-biphosphate, and a Magnesium ion. **(d)** Close-up view of one CbbL monomer, with the difference map calculated between the 3D model obtained from the map shown in (a), and the map obtained with applying C1 symmetry. A continuous density is present on one surface, exposed at the surface of the RuBisCO complex.

In addition, we observed in the map some unattributed, diffuse density at the lower contour level at the surface of the complex. This suggests that, for some of the particles, other proteins are bound in this location; however, the density is blurred, likely because of low occupancy and/or averaging during image processing. To verify this hypothesis, we performed a refinement of the RuBisCO complex without imposing any symmetry, leading to a second map at 3.8 Å resolution (Figure S2d, Table S2). Here, we observed clear continuous density that cannot be attributed to RuBisCO, on a specific surface of the protein (Figure 3d). This region of the map is at a much lower resolution, and did not allow us to identify what protein this might be based on the density alone. Nonetheless, this finding provides evidence that there are other proteins bound to RuBisCO in this region, which likely originates from the broken carboxysomes together with RuBisCO, such as CA and CsoS2 ^41^. Further investigation is required to determine which carboxysomal protein the density represents.

### Cryo-EM analysis of the α-carboxysome shell

The current structural information on α-carboxysomes is limited to low-resolution tomography data ^42,43,45,50,51^. We therefore attempted to use single-particle cryo-EM to gain insight into the overall architecture of the *Cyanobium* α-carboxysome shell. As mentioned above, the process of freezing the complex led to significant breaking, which prevented large data collection of intact carboxysomes. To address this, we froze grids immediately after purification, leading to much more intact carboxysomes. In addition, we collected data at lower magnification (Table S2), allowing a larger field of view to include more intact particles. Collectively, these strategies allowed us to collect a second dataset with an average of 2-3 intact complexes per micrographs, leading to a set of 15,545 shell particles.

Initial 2D classification of the intact carboxysome complexes was carried out (Figure S3). In these 2D classes, the cargos within the carboxysome shell are clearly ordered and organised into concentric layers, in line with the findings from previous α-carboxysome studies by electron tomography ^43,45^. 3D refinement attempts with this set of particles, without symmetry, failed to converge to interpretable models, with all the particles clustered in a small subset of angle assignments. We therefore carried out a masked 3D classification selectively for the shell (Figure S4), with icosahedral symmetry applied. This led to several classes of particles, of varying diameters from 119 nm to 123 nm (Figure S5a), demonstrating the size heterogeneity of the *Cyanobium* α-carboxysomes.

We next performed 3D refinement on the most populated class of particles, applying icosahedral symmetry with masking of the internal density. This led to a map of the carboxysome shell at ~18 Å resolution (Figure 4a, Figure S5b). At this resolution, the map is largely featureless but still allows to clearly identify the edges of the icosahedron. We also note that previous studies on synthetic BMC shells have revealed that some pseudo-hexamers form double-layered complexes that protrude from the shell surface ^21,52^. Such protrusions made of the pseudo-hexamers CsoS1D were not visible on our reconstruction, which could indicate that it is not the case for CsoS1D. Alternatively, this could indicate that CsoS1D is distributed randomly on the shell surface, and therefore double-layers are blurred out during reconstruction. A higher-resolution map, obtained without symmetry, would be required to verify this.

**Figure 4:**
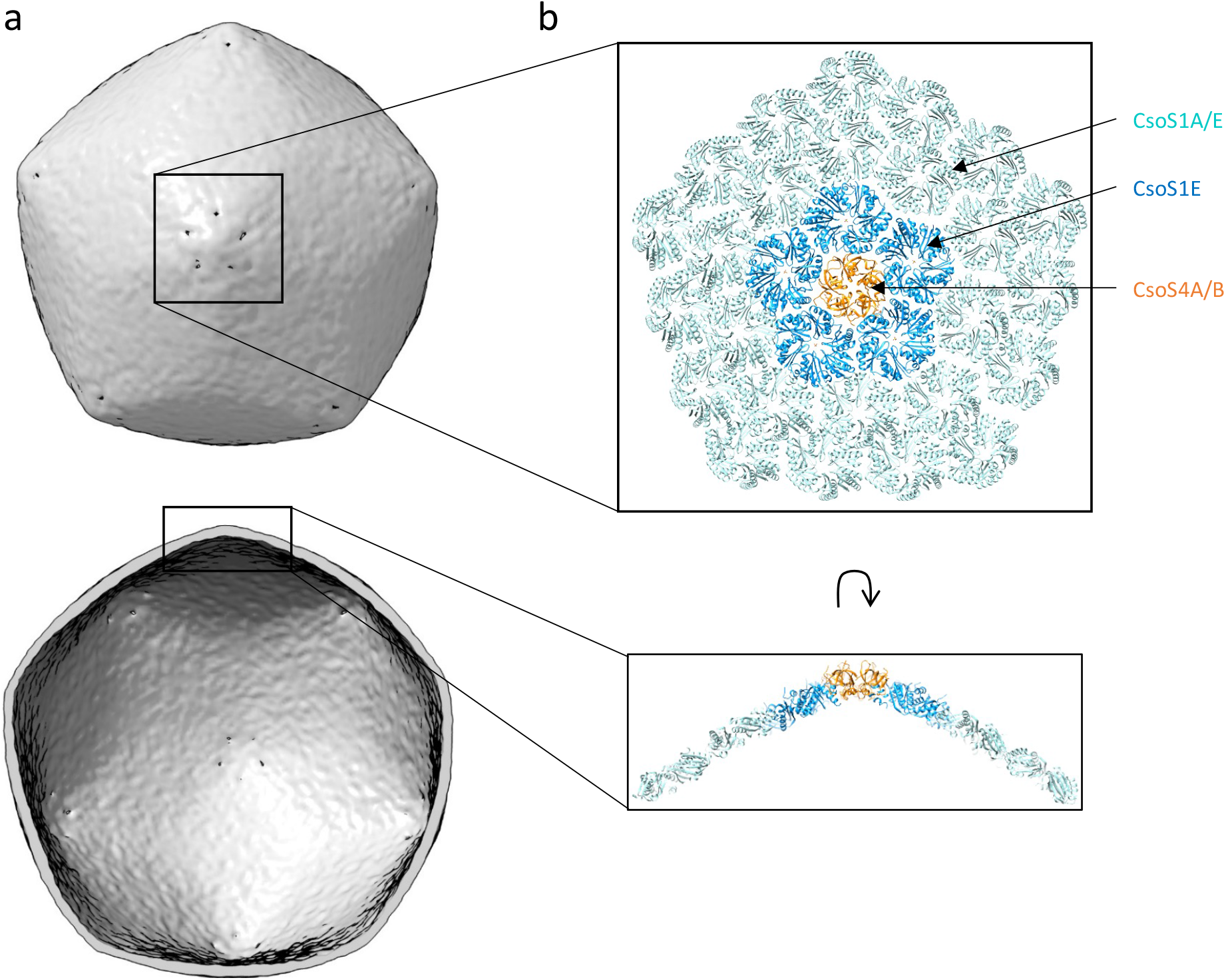
Architecture of the α-carboxysome shell. (a) Electron potential map of the carboxysome shell, to ~ 18 Å resolution. (b) Atomic model of the carboxysome shell. A single pentamer (likely a mixture of CsoS4A and CsoS4B), located at the pointy end of the shell, is shown in orange. It is surrounded by five dimers, probably consisting mostly of CsoS1E, in blue. Additional hexamer layers surround it, formed predominantly by CsoS1A, in cyan.

### Modelling of the α-carboxysome shell architecture

As indicated above, the resolution of the map of the α-carboxysome shell is not sufficient to build an atomic model *de novo*. Nonetheless, we used a hybrid approach, by combining this map with the previously elucidated structures of shell proteins and other modelling tools, to propose a structural model of the *Cyanobium* α-carboxysome shell.

Specifically, we used co-evolution analysis to determine the interactions between various shell proteins. We found a strong co-evolution correlation between CsoS1A and CsoS1E and between CsoS4A and CsoS4B (Table S3, Figures S6a, S6b). Mapping the regions with the strongest co-evolution links on the atomic models revealed that they correspond to the homo-oligomer interface (Figure S6a, S6b). The results suggest that α-carboxysome shell proteins have strong tendencies to form hetero-oligomers, i.e. hexamers formed by a combination of CsoS1A and CsoS1E, and pentamers formed of both CsoS4A and CsoS4B, as demonstrated previously in β-carboxysomes ^53,54^.

In addition, we observed a strong co-evolution correlation between CsoS1E and both CsoS4A and CsoS4B. In contrast, the correlation between CsoS1A and CsoS4A/B was very limited (Table S3). This suggests that the interaction between hexamers and pentamers is formed specifically by CsoS1E, forming the first layer of hexamers around pentamers, while CsoS1A forms predominantly hexamer-hexamer interactions. Using the hexamer and pentamer orientation derived from the previous structure of a synthetic β-carboxysome shell ^17^, the low-resolution map of the *Cyanobium* α-carboxysome shell (Figure 4a), and the co-evolution data (Table S3, Figure S6), we built an atomic model of the intact α-carboxysome shell (Figure 4b, Movie S1). In this model, the α-carboxysome shell is comprised of 12 pentamers and approximately 540 hexamers. As indicated above, there is a large variation in the dimensions of the shell, which likely corresponds to the variations in hexamer numbers. Further structural analysis, using a much larger number of intact α-carboxysome particles, is required to verify this interpretation.

Intriguingly, we observed a very limited co-evolution correlation between CsoS1D and any other shell proteins. This was likely due to its low abundance in the shell ^55^, in agreement with SDS-PAGE and mass spectrometry analysis (Figure 2, Table S1), as well as the potentially random localisation of CsoS1D in the shell facets. As such, CsoS1D is not included in this structural model. However, this protein was explicitly present within the α-carboxysome (Table S1). The role and position of CsoS1D within the shell merit further characterization.

### Internal arrangement of enzymes within the α-carboxysome

To further characterize the internal organisation of the α-carboxysomes, we carried out masked 3D refinement on the internal density (Figure S4). We initially attempted reconstructions using a range of symmetries (Figure S7); however, in most cases, this led to the blurring and distortion of features in the obtained maps. Subsequently, we applied masked three-dimensional icosahedral refinements of individual rings of densities observed within the carboxysomes. These yielded reconstructions with continuous density for each layer, which we termed the outmost, middle, inner, and core layers, respectively (Figure S8). Notably, all these layers are of a thickness that is similar to the height of RuBisCO (~ 10 nm), and possess discernible features that are suitable to its shape. We note, however, that features with 3-fold and 5-fold symmetry are present in this map, but are likely artifacts of the imposed symmetry. The thickness of each layer, and the presence of features that is compatible with RuBisCO, allowed us to manually place individual complexes in the corresponding density, leading to an atomic model of its internal organization within the carboxysome (Figure 5a, Movie S2). In this model, RuBisCO forms concentric layers, and we were able to fit ~300 RuBisCO within the internal density (4 in the core layer, 32 in the inner layer, 72 in the middle layer, and 192 in the outmost layer), roughly comparable with previous estimates ^44^. Particularly in the middle and outermost layers, gaps with thinner densities are present between RuBisCO molecules, which were not accounted for in our model. It is likely that these gaps accommodate CsoS2 and CA proteins; however, the intrinsically disordered structure of CsoS2 and the much smaller size of CA (compared to RuBisCO) did not permit us to model them within the densities.

**Figure 5:**
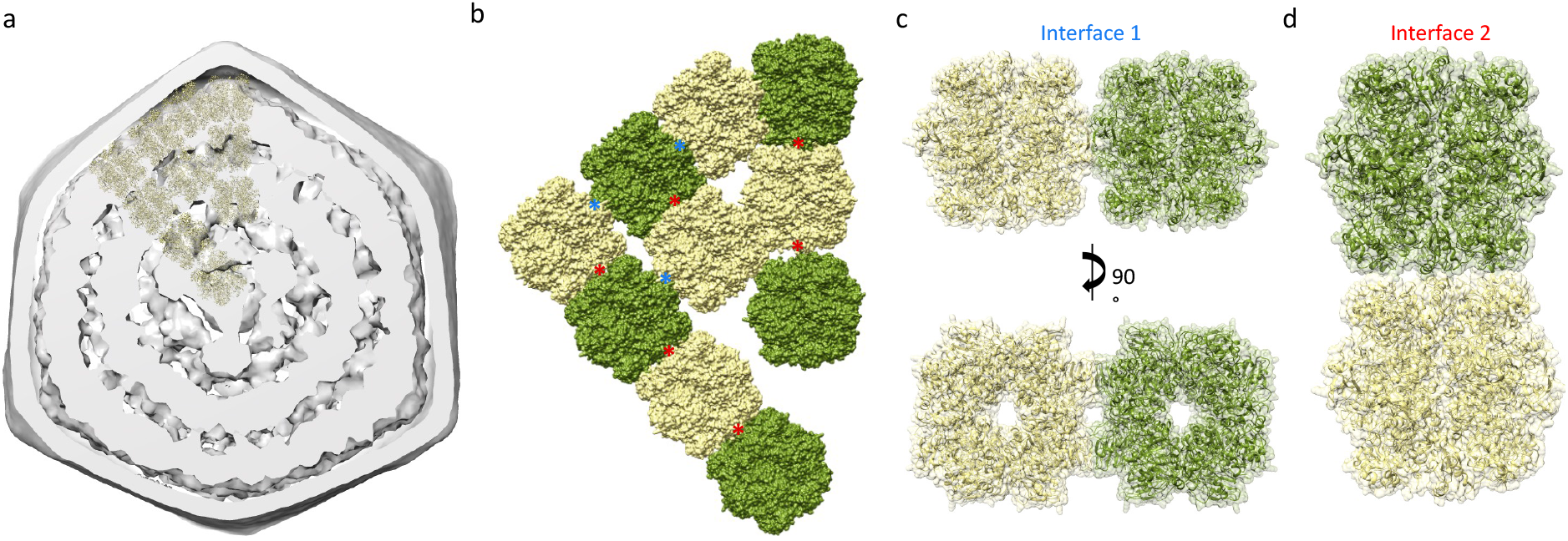
Internal arrangement of proteins within the α-carboxysome. **(a)** Slab section of the α-carboxysome electron potential map, with ten RuBisCO complexes fitted in the internal density, in cartoon representation. The height of the complex fits the with and features of the internal density. **(b)** Surface representation of ten RuBisCO complexes from the internal organization model, in surface representation, and colored alternatively in yellow and green. Two distinct inter-RuBisCO interfaces are present, indicated with a red and blue star, respectively. (c), (d) Cartoon representation of two adjacent RuBisCO molecules forming the lateral (c) and longitudinal (d) interfaces, shown in cartoon through the transparent surface. The lateral contacts occur through loops in the CbbL subunit, while the longitudinal contacts are mediated by two helices in CbbL and CbbS.

Our model of the α-carboxysomal interior organisation shows two RuBisCO interfaces (Figure 5b). The first interface corresponds to the contacts between RuBisCO proteins within the same layer, and involves interactions on the lateral side of RuBisCO (figure 5c). This interaction is presumably mediated via contacts in the variable loop region of the large subunit CbbL where CsoS2 N-terminus binds ^41^ to, which awaits further validation. A second interface is formed by the interaction between RuBisCO proteins across the concentric layers, in a top-to-bottom configuration (Figure 5b). In this case, the contacts appear to be largely mediated by two helices in the small subunit CbbS, although again the limited resolution does not allow us to unambiguously resolve this. We note, however, that diffuse density was observed in the corresponding region of the C1-derived RuBisCO map for both interfaces (see above, Figure S9), indicating the formation of residual contacts in these regions within the spilled particles.

## Discussion

In this study, we present the first single-particle cryo-EM analysis of an intact α-carboxysome, purified from endogenous sources. Notably, we report the structure of its RuBisCO to 2.9 Å, and observe the presence of unattributed densities on one side, suggesting that another protein is bound to some of the complexes. Using multistep classification, we obtained low-resolution maps of the icosahedral shell and its internal cargo organization, which allows us to propose an atomic model for their respective architecture, through integrative modelling. Collectively, this work provides insights into the architecture of BMCs and their interior organization.

We chose the *Cyanobium* α-carboxysome as a model system in this study, because its structure appears relatively more homogeneous, as demonstrated in our results (Figure S1) and previous studies ^30,49^, compared to other BMCs studied ^42–48,56^. Our results demonstrate that the *Cyanobium* α-carboxysomes exhibit an icosahedral symmetry, albeit variable in shape and size, ranging from 119 to 123 nm in diameter (Figure 2c, S5). It confirms the common icosahedral architecture of carboxysomes in different species, as observed previously ^43–45^. The model of the internal RuBisCO organization within the α-carboxysome highlights four concentric layers of cargo enzymes and two main forms (side-by-side and top-to-bottom) of RuBisCO-RuBisCO interfaces (Figure 5, S8). In contrast, recent work using cryo-electron tomography showed that in a distinct α-carboxysome from *H. neapolitanus*, RuBisCO form filaments instead of concentric layers ^51^. Nonetheless, in those filaments, the interface is highly similar to one of the interfaces identified in our model. This strongly suggests that despite the diversity of α-carboxysome species, this top-to-bottom interaction is likely a conserved feature of RuBisCO-RuBisCO association. This conserved interaction is reminiscent of the recent discovery that many metabolic enzymes, such as CTP (cytidine triphosphate) synthase and IMPDH (inosine-5’-monophosphate dehydrogenase), are able to form higher-order assemblies to regulate their activities ^57^. Whether the RuBisCO assembly patterns inside the carboxysome could modulate RuBisCO activity merits further investigation. Moreover, it is likely these filaments aid in the assembly and encapsulation of the shell in collusion with CsoS2. In comparison, Rubisco enzymes form paracrystalline arrays and exhibit relatively denser packing within the β-carboxysome ^47,48^. The discrepancy in the internal organisation and copy numbers of RuBisCO within α- and β-carboxysomes shed light on their different assembly pathways and encapsulation mechanisms.

The low resolution of the α-carboxysome map reported here, is partly due to the intrinsic heterogeneity and structural plasticity of natural carboxysome structures and internal RuBisCO packing. Given the dynamic and fast assembly, the BMC structures are morphologically heterogeneous and vary in size and shape in their native host cells ^2^. It has also been revealed that the abundance of individual proteins in the β-carboxysome and the size of β-carboxysomes in cyanobacteria could be dynamically regulated in response to changing growth conditions ^58^. Moreover, the β-carboxysome shell appeared to be mechanically softer than virus capsids, highlighting the flexible nature of the shell architecture ^48^. The structural plasticity of BMCs also occurred in protein-protein interactions, such as dynamic self-assembly and correlation between shell protein paralogs to form specific protein assemblies and hetero-oligomers in BMCs ^16,53,54,59^. Consistently, our co-evolution analysis suggests that CsoS1A, CsoS1E, CsoS4A and CsoS4B may form specific assemblies, in which CsoS4A and CsoS4B pentamers sit at the shell vertices, surrounded by CsoS1E proteins which then interact with CsoS1A hexamers (Figure 4b). It also suggests that the α-carboxysome shell paralogs CsoS1A and CsoS1E, as well as CsoS4A and CsoS4B, are prone to form hetero-oligomers (Table S3, Figure S6), as characterized in β-carboxysomes, which could function as a general mechanism for governing passage of metabolites across the carboxysome shell. These flexible interactions may play vital roles in BMC shell assembly and permeability.

The power of single-particle cryo-EM should allow obtaining the structure of an intact carboxysome to near-atomic resolution. However, there remains multiple practical challenges for this^60^. Because of the structural heterogeneity mentioned above, a very large number of particles will be required; due to the distinct symmetry between the shell and internal layers, ideally no symmetry would be applied, again increasing the number of particles required for structure determination. Additionally, the size of the complex necessitates to collect data with a large field of view, both limiting the attainable resolution and the throughput of data collection. Finally, the size of the data, and of the particles used for reconstruction, presents a challenge for the data processing. Nonetheless, with the most recent wide-field direct-electron cameras ^61^, and with improved automation in data acquisition, this will likely be obtainable in the future. We also emphasize that tomography approaches will likely also provide important new insights on the structural diversity of these complexes. While likely not providing high-resolution structural information, tomography will notably be key to identify and characterize assembly intermediates. Nonetheless, data collection for tomography is even more challenging than for single-particle analysis, and its low-throughput likely remains a limiting factor.

Recently, extraordinary advances have been made in the acquisition of high-resolution characterization of synthetic BMC minishells ^19–21,52,62^. These synthetic shells, with minimal components, exhibit more homogeneous structures and lack any of the internal enzymes, thereby facilitating the alignment of the particles. In contrast, our study on the intact α-carboxysome structure provides insights into the carboxysome assembly as well as the diversity of BMC architectures and protein compositions. Further characterizations are expected to address how CsoS2 assists with the association of the outer layer of Rubisco and shell proteins, how CsoS1D and CA are organised within the native α-carboxysome, and how the internal packing of RuBisCO enzymes is physiologically regulated.

## Materials and Methods

### Cyanobacterial strain growth and carboxysome purification

*Cyanobium* sp. PCC 7001 (Pasteur Culture Collection of Cyanobacteria, PCC) cells were grown in 4 L of BG-11 medium under constant illumination at 30°C with constant stirring and bubbling with air. Carboxysomes were purified as described previously with modifications. Cells were collected by centrifugation (6000 g, 10 min) and resuspended in TEB buffer (5 mM Tris-HCL, pH 8.0, 1 mM EDTA, 20 mM NaHCO_3_) with additional 0.55 M mannitol and 60 kU rLysozyme (Sigma-Aldrich, United States). Cells were then incubated overnight (20 h) with gentle shaking at 30°C in the dark, and were collected via centrifugation (6000 g, 10 min). Cells were placed on ice and resuspended in 20 mL ice-cold TEB containing an additional 5 mL 1 μm Silicone disruption beads. Cells were broken via bead beating for 8 mins in one-minute intervals of vortex, and 1 min on ice. Broken cells were separated from the beads, and the total resuspension volume was increased to 40 mL with TEB buffer containing an additional 4% IGEPAL CA-630 (Sigma-Aldrich, United States) were mixed on a rotating shaker overnight at 4°C. Unbroken cells were pelleted via centrifugation at 3,000 g for 5 mins, and the supernatant was centrifuged at 40,000 g for 20 mins. The pellet was then resuspended in 40 mL TEMB containing 4% IGEPAL CA-630 and centrifuged again at 40,000 x g for 20 mins. The resulting pellet was then resuspended in 2 mL TEB + 10mM MgCl_2_ (TEMB) (5 mM Tris-HCL, pH 8.0, 1 mM EDTA, 10 mM MgCl_2_, 20 mM NaHCO_3_) and centrifuged at 5000 x g for 5 mins before loading onto a 20-60% (v/v) sucrose gradient in TEMB buffer. Gradients were then centrifuged at 105,000 g for 60 mins at 4°C; the milky band at the 40%-50% interface was collected, diluted in 10 mL TEMB buffer and centrifuged again at 105,000 g for 60 mins. The final carboxysome pellet was then resuspended in 150 μL TEMB for the following structural and biochemical analysis.

### SDS-PAGE and immunoblot analysis

Isolated carboxysomes were diluted to 5 mg mL^-1^ and denatured using 4X Bromophenol blue buffer (Fisher Scientific, United States). The samples were heated at 95°C for 10 mins, and insoluble debris was pelleted via short spin. Approximately 50 μg proteins were loaded onto 15% (v/v) denaturing SDS-PAGE gels and stained using Coomassie Brilliant Blue G-250 (ThermoFisher Scientific, UK). Immunoblot analyses were performed using anti-RbcL (1:10,000 diution, Agrisera, AS03 037, Sweden), anti-CsoS1 from *H. neapolitanus* (1:5000 dilution, Agrisera, AS14 2760, Sweden), and horseradish peroxidase-conjugated goat antirabbit immunoglobulin G secondary antibody (1:10,000 dilution, Agrisera AS101461, Sweden). Images were taken using a Quant LAS 4000 platform (GE Healthcare Life Sciences, USA)

### RuBisCO assay

RuBisCO activities of isolated carboxysomes were determined as described previously with minor modifications ^26,32,37^. Isolated α-carboxysomes were diluted to 0.5 mg mL^-1^ in (100 mM EPPS, pH 8.0; 20 mM MgCl_2_) and 5 μL was added to scintillation vials containing NaH^14^CO_3_ with a range of concentrations (1.5-48 mM). and incubated at 37 °C for 2 mins before the addition of D-ribulose 1,5-bisphosphate sodium salt hydrate (RuBP, Sigma Aldrich, US) final concentration 0.04 mM. The reaction was carried out for 5 mins before being terminated by adding 2:1 by volume 10% formic acid. Samples were dried for at least 30 mins at 95 °C to remove unfixed ^14^C before re-suspending the fixed ^14^C pellets with ultra-pure water and adding 2 mL of scintillation cocktail (Ultima Gold XR, PerkinElmer, US). Radioactivity measurements were then performed using a scintillation counter (Tri-Carb, PerkinElmer, US). Raw readings were used to calculate the amount of fixed ^14^C, and then converted to the total carbon fixation rates. RuBisCO activity. Data are presented as mean ± standard deviation (SD) based on three biological replicates isolated from independent culture batches, and were analyzed using OriginPro 2020b (OriginLab, Massachusetts, USA).

### Mass spectrometry analysis

The isolated α-carboxysome samples were washed with PBS buffer. Rapigest was added to a final concentration of 0.05% (w/v) into the sample for 10-min incubation at 80°C. The sample was then reduced with dithiothreitol (3 mM, final concentration) for 10 mins at 60°C, alkylated with iodoacetamide (9 mM, final concentration) for 30 min at room temperature in the dark, followed by digestion with trypsin at 37°C overnight. Digestion was terminated with 1 μL of trifluoroacetic acid (TFA). Data-dependent LC-MS/MS analysis was conducted on a QExactive quadrupole-Orbitrap mass spectrometer coupled to a Dionex Ultimate 3000 RSLC nano-liquid chromatograph (Hemel Hempstead, UK). A 2 μL sample digest was loaded onto a trapping column (Acclaim PepMap 100 C18, 75 μm × 2 cm, 3 μm packing material, 100 Å) in 0.1% TFA, 2% acetonitrile H_2_O, and set in line with the analytical column (EASY-Spray PepMap RSLC C18, 75 μm × 50 cm, 2 μm packing material, 100 Å). Peptides were eluted using a linear gradient of 96.2% buffer A (0.1% formic acid):3.8% buffer B (0.1% formic acid in water:acetonitrile 80:20, v/v) to 50% buffer A:50% buffer B over 30 mins at 300 nL min^-1^. The mass spectrometry analysis was operated in DDA mode with survey scans between *m*/*z* 300-2000 acquired at a mass resolution of 70,000 (FWHM) at *m*/*z* 200. The maximum injection time was 250 ms, and the automatic gain control was set to 1e^6^. Fragmentation of the peptides was performed by higher-energy collisional dissociation using a normalized collision energy of 30%. Dynamic exclusion of *m*/*z* values to prevent repeated fragmentation of the same peptide was used with an exclusion time of 20 seconds.

The raw data file was imported into Progenesis QI for Proteomics (Version 3.0 Nonlinear Dynamics, Newcastle upon Tyne UK, a Waters Company). Peak picking parameters were applied with sensitivity set to maximum and features with charges of 2^+^ to 7^+^ were retained. A Mascot Generic File, created by Progenesis, was searched against the *Cyanobium* sp. PCC 7001 database from UniProt (UP000003950, 2762 proteins) with the sequence of yeast enolase (UniProt: P00924) added. Trypsin was specified as the protease with one missed cleavage allowed and with fixed carbamidomethyl modification for cysteine and variable oxidation modification for methionine. A precursor mass tolerance of 10 ppm and a fragment ion mass tolerance of 0.01 Da were applied. The results were then filtered to obtain a peptide false discovery rate of 1%. Protein quantification was calculated using Hi3 methodology using yeast enolase (50 fmol μL^-1^) as a standard protein.

### Thin-section electron microscopy

Cyanobacterial cell cultures were pelleted by centrifugation (6,000 g, 10 min) and processed for thin section using a Pelco BioWave Pro laboratory microwave system. The cells are first fixed with 2.5% glutaraldehyde in 0.1 M sodium cacodylate buffer at pH 7.2 using two steps of 100W. After agarose embedding, samples were then stained with 2% osmium tetroxide and 3% Potassium Ferrocyanide using three steps of 100W. The osmium stain was set using 1% thiocarbohydrazide and 2% osmium tetroxide. The samples were stained with 2% uranyl acetate, prior to dehydration by increasing alcohol concentrations (from 30 to 100%) and resin embedding. Thin sections of 70 nm were cut with a diamond knife and poststained with 3% lead citrate.

### Negative-stain TEM grid preparation and screening

Isolated α-carboxysome samples were immobilized onto the glow-discharged grids and then were stained with 2% uranyl acetate. EM imaging was conducted using an FEI Tecnai G2 Spirit BioTWIN transmission electron microscope equipped with a Gatan Rio 16 camera.

### Cryo-EM grid preparation and data collection

For the structural characterisation of RuBisCO, 3 μL aliquots of purified α-carboxysomes at a concentration of ~1 mg mL^-1^ were applied to Graphene Oxide coated, 300 mesh, 2/2 μm hole/spacing, holey carbon grids (EMR). A Leica EM GP Automatic Plunge Freezer (Leica) was used to plunge freeze the sample, blotting for 3-6 s. Cryo-EM data was collected with a 300 kV Titan Krios TEM, equipped with a Falcon 3 direct electron detector (Thermo Fisher) operated in linear mode. 4593 micrographs were collected using the EPU software (Thermo Fisher) with a pixel size of 1.11 Å pix^-1^, a total dose rate of 30 e^-^ Å^-2^, and 44 fractions per micrograph. The defocus range was −0.5 to −1.5 μm.

For structural characterisation of the intact α-carboxysome complex, 3 μL aliquots of purified sample at a concentration of 3 mg mL^-1^ were applied to Graphene Oxide coated grids, 300 mesh, 2/2 μm hole/spacing, holey carbon grids (EMR). A Leica EM GP Automatic Plunge Freezer (Leica) was used to plunge freeze, blotting for 6 s. Cryo-EM data were collected with a 300 kV Titan Krios TEM with a Falcon 3 direct electron detector (FEI) operated in counting mode. 5429 micrographs were collected using EPU software (Thermo Fisher) with a pixel size of 2.23 Å pix^-1^ with a total dose rate of 29.7 e^-^ Å^-2^ with 33 frames per micrograph. The defocus range was −1.0 to −2.2 μm.

### Cryo-EM data processing

All the cryo-EM data processing steps were carried out in CryoSPARC ^63^v.3.1.0.. For the RuBisCO structure, automated particle picking was initially used, leading to a dataset of ~ 2,800,000 particles. 2D classification was employed to select particles that clearly correspond to RuBisCo, leading to a final set of 131,356 particles. 3D refinement was performed with these, with D4 symmetry, converging to a map at 2.87 Å resolution. The same set of particles was also refined without symmetry imposed, leading to a second map at 3.79 Å resolution.

For intact carboxysomes, 131 particles were manually picked from selected micrographs to generate 2D classes subsequently used for template picking for the entire dataset. A total of 15545 particles were picked and extracted using a 700×700 pixels box. After multiple rounds of 2D classification 8701 particles from the best 2D classes were selected and used to generate an initial model. Particles were downsampled to a box size of 168×168 pixels for 3D classifications and reconstructions. A reconstruction of the entire carboxysome was generated in I symmetry. Masked classifications of the shell were carried out with C1 symmetry to give a reconstruction at 19 Å resolution. Heterogeneous refinements of the carboxysome shell used for model building were carried out with I symmetry to give reconstructions of ~18 Å.

### Modelling and co-evolution analysis

Atomic models of the CsoS1A and CsoS1E hexamers, the CsoS4A and CsoS4B pentamers, and the CsoS1D trimer were generated with AlphaFold^64^. The co-evolution analyses were performed using the RaptorX server^65^, with contact probabilities > 0.5 considered to be significant.

To build the *Cyanobium* sp. PCC 7001 RuBisCO structure, an initial atomic model was built for both CbbL and CbbS with AlphaFold, and 8 copies of each were placed at their respective location on the EM map. The coordinates for the substrate and Mg ion were added manually, and the termini without visible density were deleted. The model was then subjected to real-space refinement in Phenix^66^.

The difference map was calculated by first generating a volume of the RuBisCO structure, and then subtracting this volume from the C1 reconstruction, in ChimeraX ^67^.

To generate the atomic model of the shell, a CsoS4a pentamer was placed in one corner of the map icosahedron, using the orientation reported previously in the structure of the β-carboxysome synthetic shell ^21^ to determine the outward face. Five copies of the CsoS1E hexamer were placed around it, again using the β-carboxysome structure to determine the outward face, and the interface was optimized manually by fitting to the map in Chimera ^68^. Additional copies of the CsoS1A hexamers were next placed manually, forming two additional continuous layers around. Further extension of the model with additional CsoS1A hexamers could not form continuous layers, and included significant gaps; nonetheless, the number of hexamers required to complete the map could be estimated by placing as many as possible in the volume without any significant clashes.

For the internal density, copies of the *Cyanobium* sp. PCC 7001 RuBisCO structure were placed in regions of the map of the different shells, and fitted manually in Chimera. If major clashes were observed between adjacent molecules, that with the less optimal fit to the density was removed.

All structural figures were generated in either PyMol ^69^, Chimera, or ChimeraX.

## Acknowledgements

This work was supported by the National Natural Science Foundation of China (32070109), the National Key R&D Program of China (2021YFA0909600), the Biotechnology and Biological Sciences Research Council Grant (BB/R019061/2 to JRCB, BB/M024202/1, BB/V009729/1, BB/R003890/1 to LNL), the Royal Society (RGF\EA\180233, URF\R\180030, RGF\EA\181061 to LNL), the Leverhulme Trust (RPG-2021-286 to LNL). We acknowledge Diamond Light Source for access and support of the cryo-EM facilities at the UK’s national Electron Bio-imaging Centre (eBIC) (under proposal EM-19832). We thank Prof. Ian Prior and Mrs. Alison Beckett for the support of electron microscopy, and Dr. Deborah Simpson and Dr. Philip J Brownridge for mass spectrometry analysis. The University of Sheffield FoS cryo-EM facility was used for initial grid preparation and optimization. We are grateful to Dr Alex Parker for help with generating the supplementary movies.

## Author Contributions

SLE performed the cryo-EM sample preparation and data processing for the shell and internal organization, with help from DM; as well as the hybrid structural modelling, with help from JRCB. MMJA performed cell culturing, carboxysome isolation, biochemical characterization and RuBisCO assays, with help from YS and TC. GFD performed negative staining TEM sample preparation and imaging. DM and NS performed the grid preparation and data processing for the RuBisCO structure. AJB helped with cryo-EM data transfer and processing. SLE, LNL, and JRCB wrote the manuscript, with contributions from all the authors.

## Data availability

The structure of the *Cyanobium* sp. PCC 7001 RuBisCO enzyme has been deposited to the PDB (ID: 7YYO), and the corresponding 2.9 Å cryo-EM map was deposited to the EMDB (ID: 14385). The map obtained without imposing any symmetry, to 3.8 Å, was also deposited (ID: 14376). The maps of the carboxysome shell, and of each individual internal layer, have been deposited to the EMDB (ID: 14379, 14382, 14381, 14380, and 14377, respectively).

## Notes

### Competing Interest Statement

The authors have declared no competing interest.

