## Supplementary material for "Single-particle cryo-EM analysis of the shell architecture and internal organization of an intact α-carboxysome"

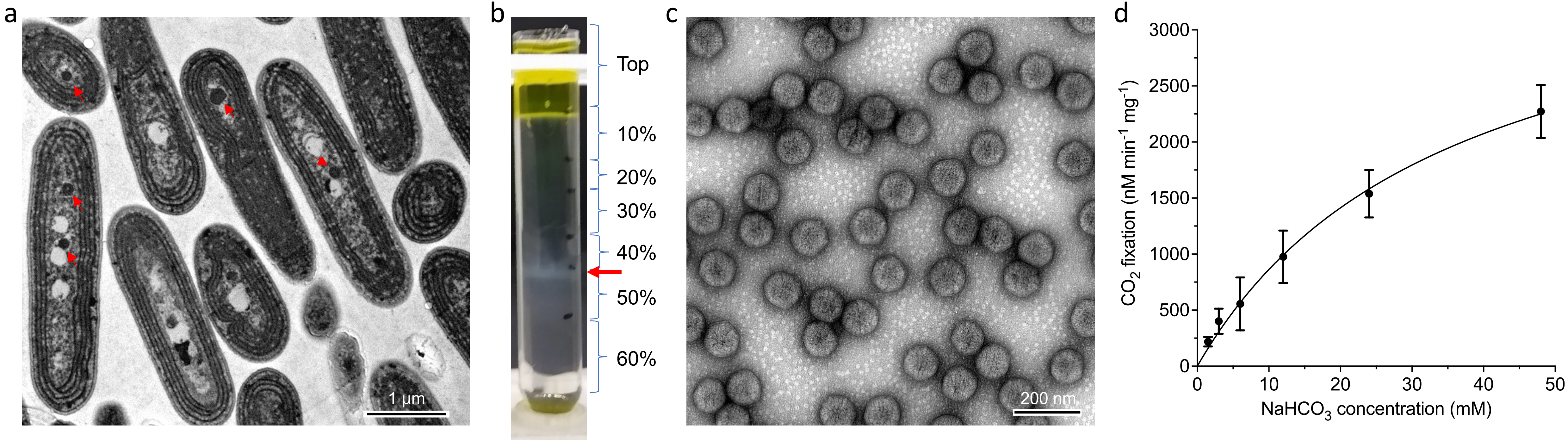


**Figure S1: Isolation and purification of the *Cyanobium* α-carboxysome**. **(a)** Thin section of cultured cells, with carboxysomes indicated with red arrows. **(b)** Sucrose gradient from the purification process. Carboxysome-enriched samples are indicated in red arrow. **(c)** Negative-stain micrograph of purified carboxysome complexes. RuBisCO molecules, presumably from broken carboxysome complexes, can be seen in the background. **(d)** RuBisCO activity assay from the purified α-carboxysomes, demonstrating that they are functional for carbon fixation.

| Protein | Score | Mass (Da) | Normalized amount (fmol) |
|---|---|---|---|
| **CbbL** | 3376.57 | 53050 | 8682.5 ± 2435.3 |
| **CsoS2** | 5646.53 | 86750 | 1288.5 ± 194.5 |
| **CSoS1A** | 1661.34 | 10628 | 9654.3 ± 192.0 |
| **CbbS** | 505.73 | 13061 | 4659.1 ± 3110.8 |
| **CsoSCA** | 1771.19 | 61331 | 118.0 ± 18.0 |
| **CsoS1E** | 831.92 | 18563 | 181.1 ± 55.6 |
| **CsoS4A** | 104.78 | 10317 | 5.4 ± 8.8 |
| **CsoS1D** | 327.88 | 26073 | 1.6 ± 0.1 |
| **CsoS4B** | 145.98 | 8894 | 1.0 ± 1.7 |
| **Bacterioferritin** | 687.17 | 17782 | 88.7 ± 49.1 |
| **HAM1** | 451.01 | 20496 | 8.6 ± 5.2 |

**Table S1: Proteomic results of isolated α-carboxysomes from *Cyanobium*.** The column of Normalized amount displays the amount of each of the carboxysomal proteins detected in isolated α-carboxysomes using mass spectrometry, normalized against the amount of the least abundant protein CsoS4B. Note: Mass spectrometry revealed the presence of bacterioferritins and purine NTP pyrophosphatases in the isolated α-carboxysome samples. Both genes encoding bacterioferritin (CPCC7001_1612) and purine NTP pyrophosphatase (CPCC7001_2175) are located within the α-carboxysome operon in the *Cyanobium* genome.

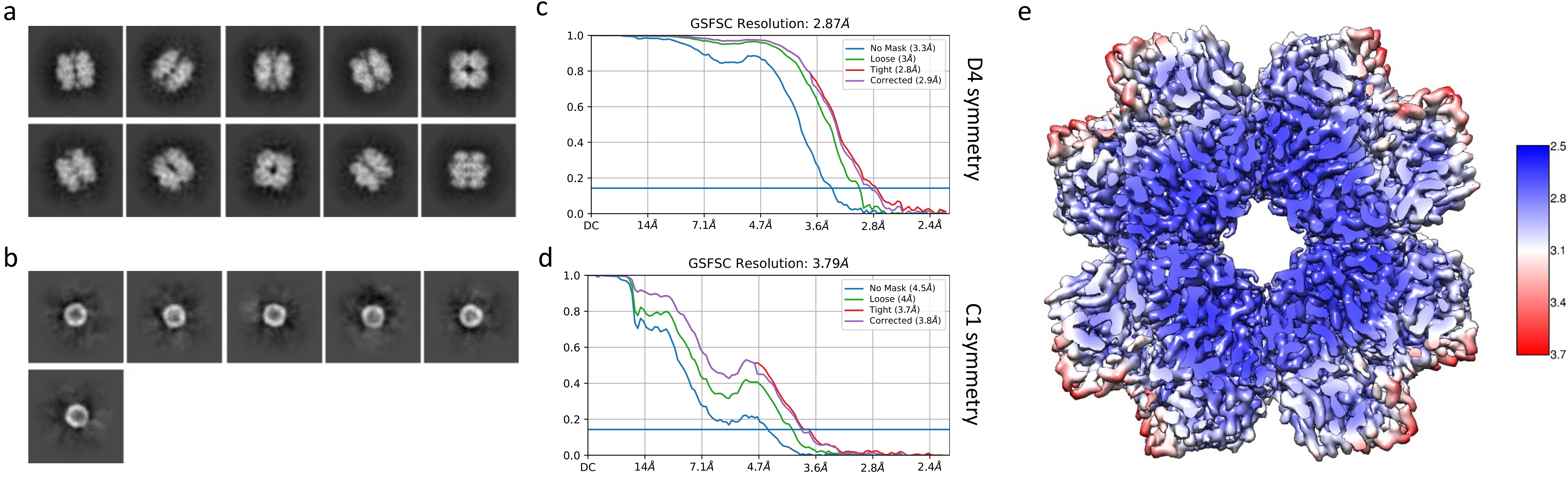


**Figure S2: Cryo-EM reconstruction of the spilled RuBisCo. (a), (b)** 2D classification of the proteins spilled from broken α-carboxysomes**.** (a) A subset of particles formed highly-ordered 2D classes, whose size and overall shape matched that of RuBisCo. (b) A second sub-set of particles were smaller, and not well resolved. **(c), (d)** Half-map FSC curves for the RuBisCo structures, with D4 symmetry (c) and with C1 symmetry (d). **(e)** RuBisCo electron potential map colored by local resolution.

| | Dataset 1 | | Dataset 2 |
|---|---|---|---|
| **Data collection** | | | |
| Voltage (kV) | 300 | | 300 |
| Exposure (e/Å$^2$) | 30 | | 29.7 |
| Fractions | 44 | | 33 |
| Defocus range ($\mu$m) | -0.5 to -1.5 | | -1.0 to -2.2 |
| Pixel size (Å pix$^{-1}$) | 1.11 | | 2.23 |
| Number of micrographs | 4593 | | 5429 |
| Initial particle number | 2,800,000 | | 15,545 |
| **Map refinement** | | | |
| Final particle number | 131,356 | 131,356 | 3,533 |
| Resolution (Å) | 2.87 | 3.79 | 18.25 |
| Symmetry | D4 | C1 | I |
| **Structure refinement** | | | |
| Non-hydrogen atoms | 35,136 | | |
| Protein residues | 4,424 | | |
| Ligands | 16 | | |
| Protein B-factor | 73.68 | | |
| Ligand B-factor | 61.35 | | |
| Bond length RMSD (Å) | 0.008 | | |
| Bond angle RMSD (º) | 0.966 | | |
| MolProbity score | 2.84 | | |
| Clash score | 12.58 | | |
| Poor rotamers | 12.83 | | |
| Ramachandran favoured (%) | 94.33 | | |
| Ramachandran allowed (%) | 5.67 | | |
| Ramachandran disallowed (%) | 0.00 | | |

**Table S2: Cryo-EM data collection and structure refinement parameters**

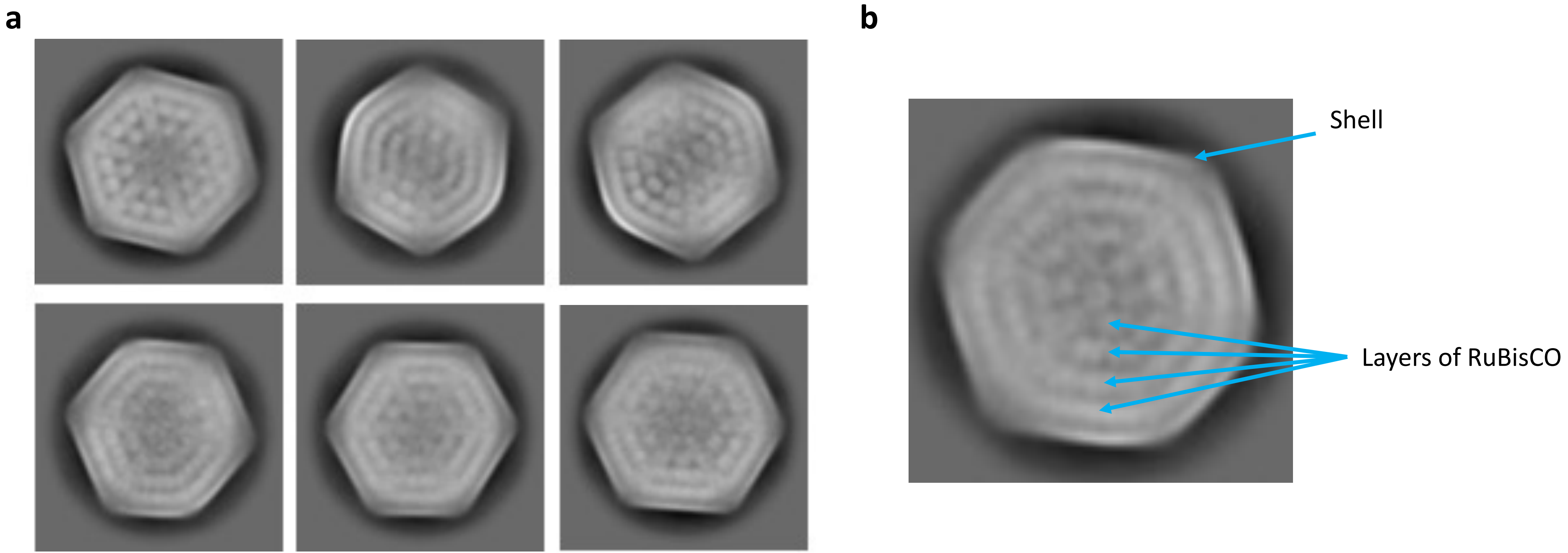


**Figure S3: 2D classification of intact α-carboxysome particles.** (a) selected 2D classes of α-carboxysome complexes from the second dataset, demonstrating a rigid organization of the internal enzymes. (b) Close-up view of one such 2D class, with the localization of the shell, and internal RuBisCO layers

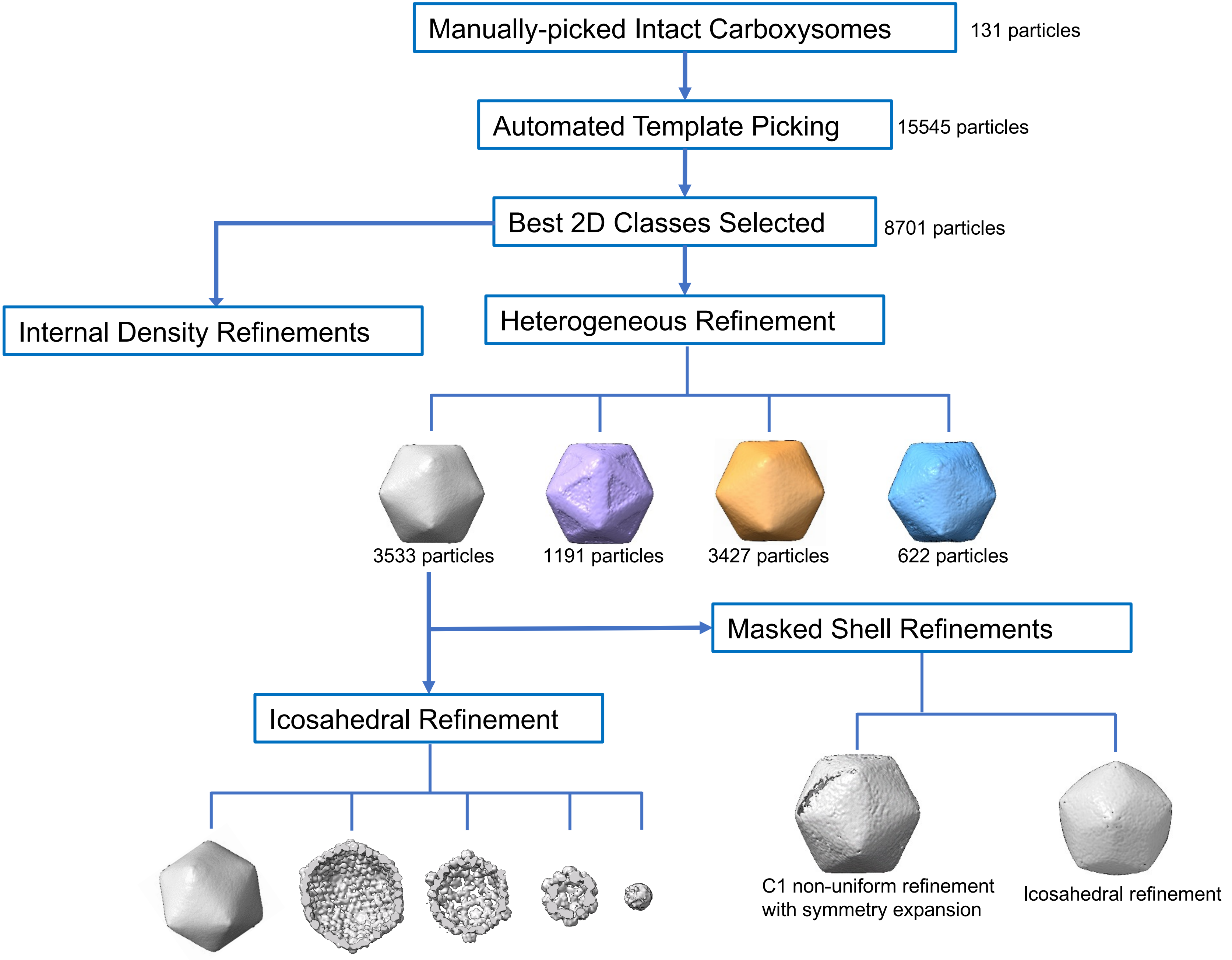


**Figure S4: α-carboxysome cryo-EM processing pipeline.** The various steps used for the processing of the 2nd dataset are indicated.

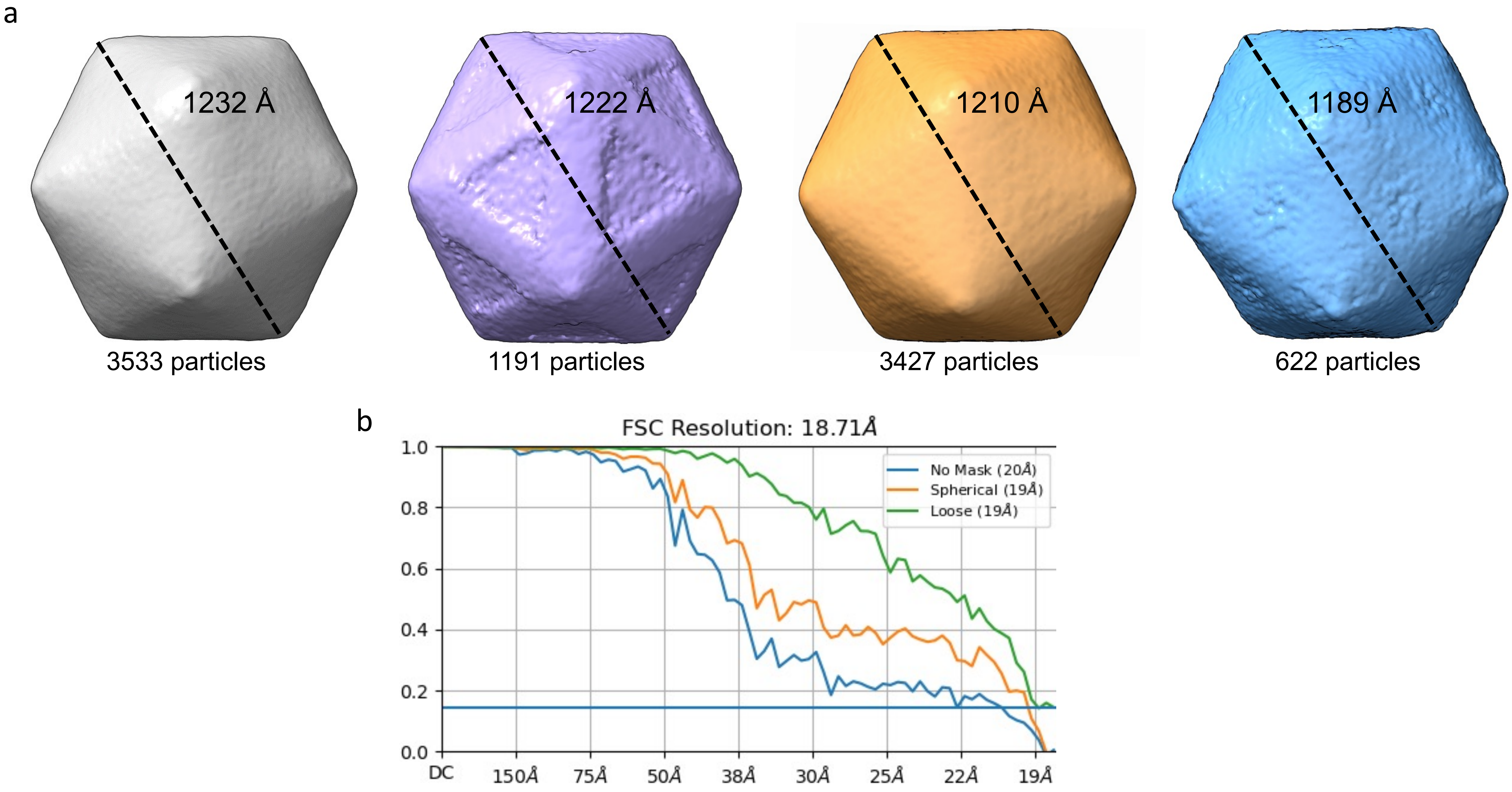


**Figure S5: Classification and refinement of the α-carboxysome shell.** (a) 3D classes of the shell particles. The for each class, the number of particles and the shell diameter are indicated. (b) FSc curve for the carboxysome shell map, with Icosahedral symmetry.

| | CsoS1A | CsoS1D | CsoS1E | CsoS4A | CsoS4B | CsoS2 |
|---|---|---|---|---|---|---|
| **CsoS1A** | / | 0 | 8 | 1 | 3 | 14 |
| **CsoS1D** | / | / | 2 | 1 | 0 | 2 |
| **CsoS1E** | / | / | / | 4 | 6 | 12 |
| **CsoS4A** | / | / | / | / | 1 | 2 |
| **CsoS4B** | / | / | / | / | / | 1 |
| **CsoS2** | / | / | / | / | / | / |

**Table S3: Co-evolution analysis of the shell proteins.** The number of co-evolving residues with a score > 0.5 for each protein pair is indicated.

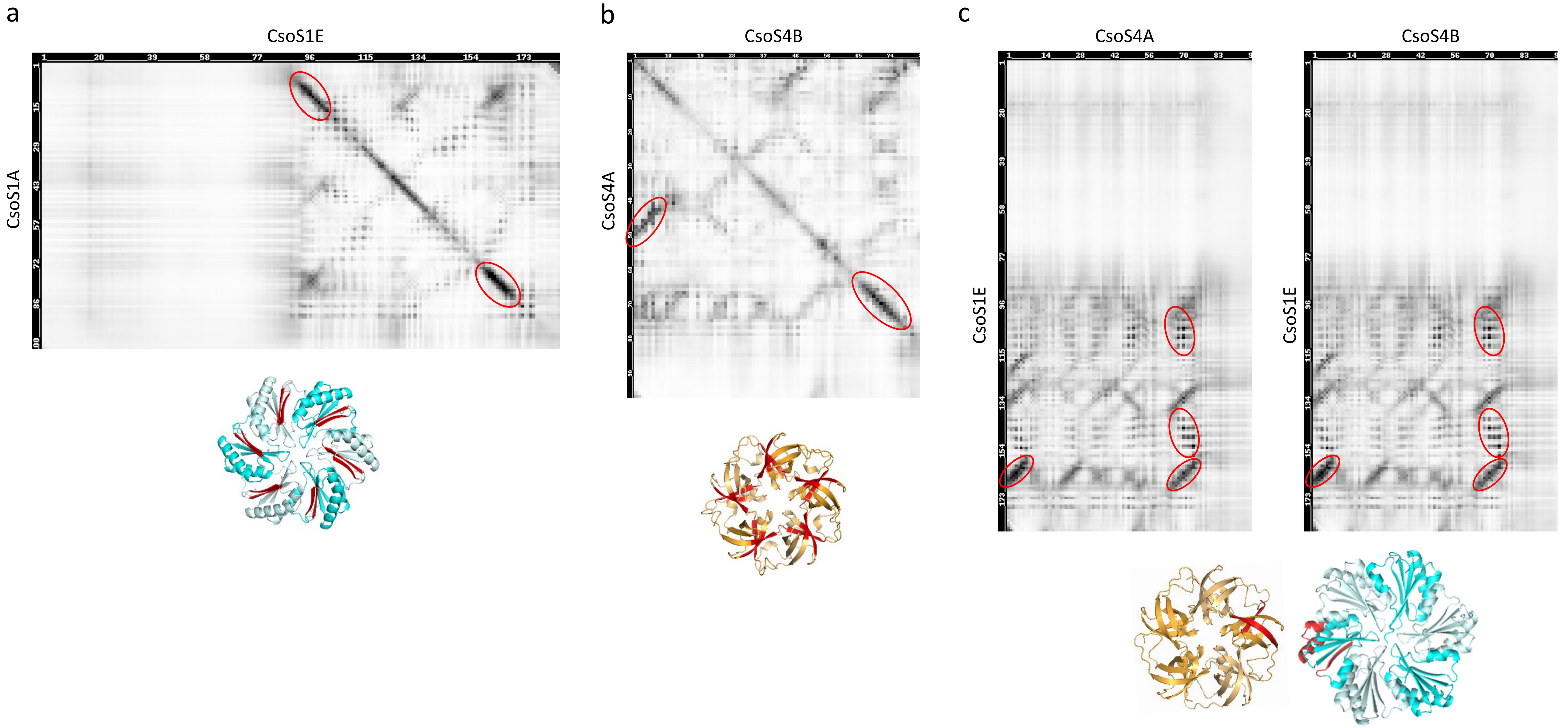


**Figure S6: Co-evolution maps of the α-carboxysome shell proteins.** For each protein pair, the co-evolution map is shown at the top, and an atomic model is at the bottom, colored as in figure 1. The residues with strong co-evolution correlation are indicated with a red circle, and colored in red in the structural model. The co-evolution analysis for CsoS1A and CsoS1E **(a)**, as well as CsoS4A and CsoA4B **(b)** strongly suggest inter-homooligomer interactions. In contrast, in the case of CsoS1E and CsoS4A/B, intra-homooligomer interactions are likely identified.

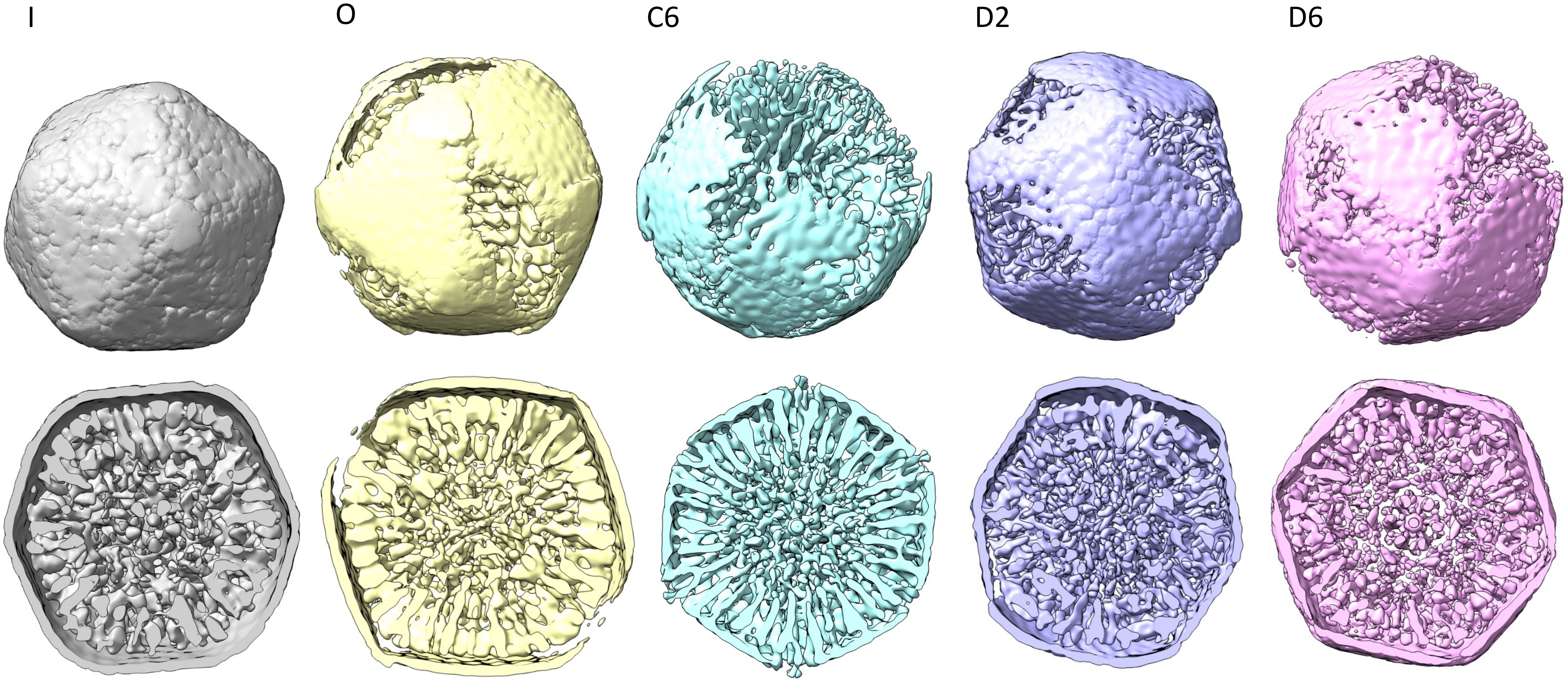


**Figure S7: 3D refinement of intact α-carboxysome particles, with a range of symmetries.** For each symmetry used, the overall map, as well as a transversal section through the density to reveal the internal layers.

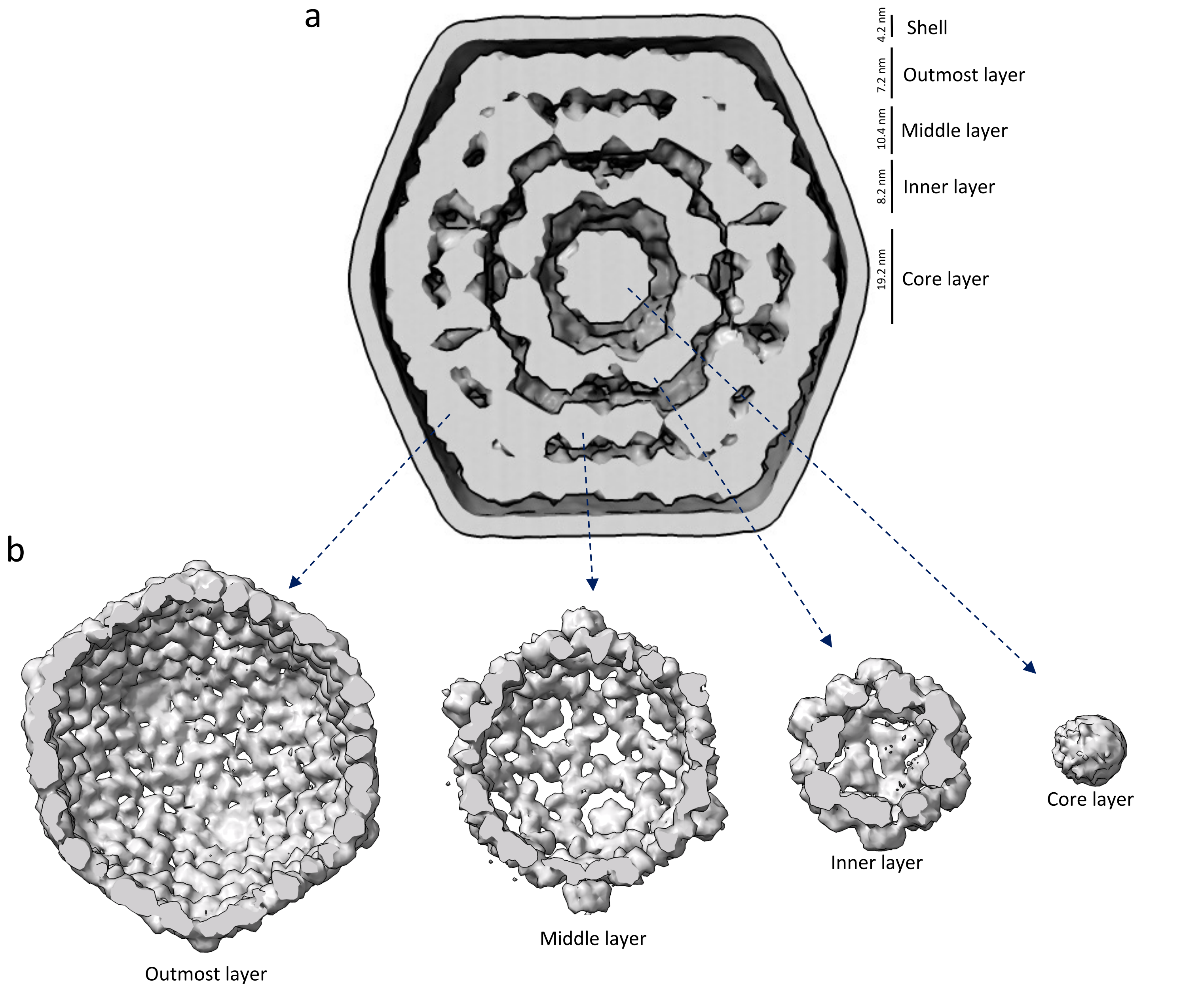


**Figure S8: Internal density of the α-carboxysome structure.** (a) Slab through the overall density, revealing the different internal layers**.** The height of each layer is indicated. (b) Individual maps for each layer, obtained from selective masked refinement of the particles used in (a).

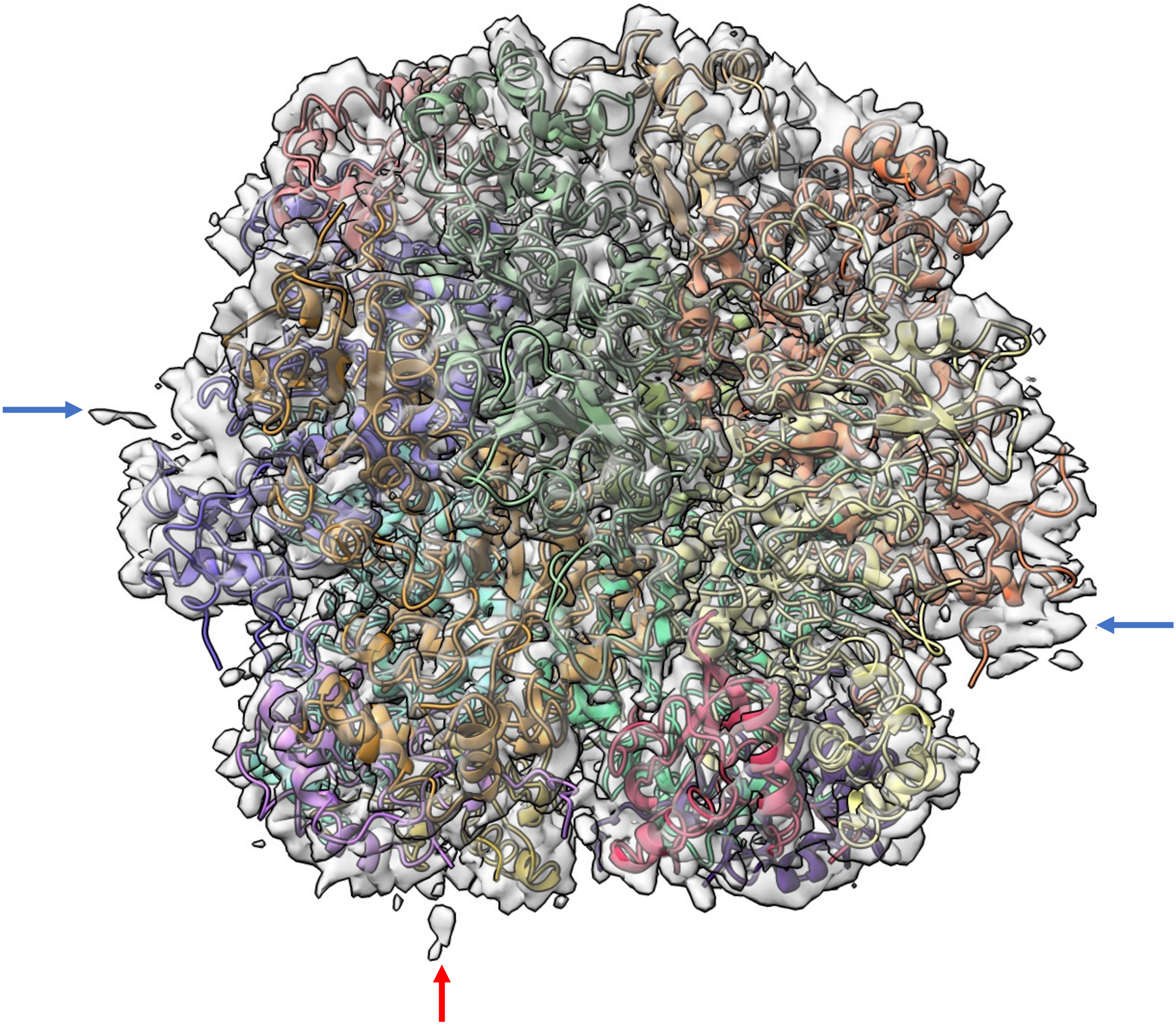

**Figure S9: Density at the RuBisCO oligomerization interface.** The Electron potential map of RuBisCO, obtained without symmetry, is shown, with the cartoon representation of the atomic model derived from the 2.9A structure visible through. Diffuse density on the edges, indicated with blue arrows, correspond to the lateral contacts in the carboxysome internal organization model, while density at the bottom, shown with a red arrow, is located in a position corresponding to the longitudinal interface.
